# The role of GATA2 in the expression of the soluble decoy receptor ST2/IL1RL1 in human and mouse mast cells

**DOI:** 10.1101/2025.02.19.638998

**Authors:** Kazuki Nagata, Kazumi Kasakura, Kenta Ishii, Naoto Ito, Mutsuko Hara, Ko Okumura, Chiharu Nishiyama

**Author notes:** Correspondence should be addressed to: Chiharu Nishiyama, Ph.D., Department of Biological Science and Technology, Faculty of Advanced Engineering, Tokyo University of Science, 6-3-1 Niijuku, Katsushika-ku, Tokyo 125-8585, Japan. These authors contributed equally to this work (Kazuki Nagata, Kazumi Kasakura).

## Abstract

The *ST2/IL1RL1* gene encodes a receptor subunit for IL-33. The *ST2/IL1RL1* gene is transcribed and translated to two splice variants, full-length ST2 (ST2L) and soluble ST2 (sST2). ST2L is expressed on cell surface as membrane bound molecule and forms a heterodimer with IL-1RAcP, which plays an important role in Th2 transduces a signaling of IL-33. In contrast, sST2 blocks the IL-33-signaling via functioning as a decoy receptor of IL-33. In this study, we analyzed the regulatory mechanisms of the gene expression of sST2 in mast cell, the major source of sST2. We found that a hematopoietic cell-specific transcription factor GATA2 is essential for sST2 expression in human mast cells (a cell line and primary cells) and mouse mast cells. Various assays including ChIP assay, reporter assay, and 3C assay revealed that GATA2 binding to the distal promoter of the *IL1RL1* gene transcriptionally activates proximal promoter-driven expression of sST2 through chromosomal conformation.

---

The *ST2/IL1RL1* gene encodes a receptor subunit for IL-33. It is transcribed and translated into two splice variants, full-length ST2 (ST2L) and soluble ST2 (sST2). ST2L is expressed on the cell surface as a membrane bound molecule and forms a heterodimer with IL-1RAcP, which plays an important role in transducing IL-33 signals in Th2. In contrast, sST2 blocks IL-33 signaling by functioning as a decoy receptor of IL-33. Although we previously reported the transcriptional regulation of ST2L in human mast cell (MC) ^1^, a major target of IL-33, the mechanisms underlying the expression of sST2 remain unclear. Therefore, we initially examined the expression of sST2 and ST2L in mouse bone marrow-derived MCs (BMMCs). The results obtained showed that a Ca^2+^ ionophore- or IgE-mediated stimulation released the sST2 protein from BMMCs (**Fig. 1A**), whereas surface levels of ST2L on BMMCs were down-regulated by a treatment with A23187 (**Fig. 1B**). Semi-quantitative PCR of *Il1Rl1* mRNA revealed that IgE-dependent activation accelerated the proximal promoter-driven transcription of the *Il1Rl1* gene and increased sST2 mRNA levels (**Fig. 1C**). The stimulation-induced increase in sST2 release (**Fig. 1D**) and decrease in surface ST2L (**Fig. 1E**) were also observed in a the human MC, LAD2 ^2^. The distal and proximal promoters were both activated for the transcription of sST2 in LAD2, while the sST2 transcript was driven by the proximal promoter and ST2L was not expressed in human keratinocytes (**Fig. 1F**). The structures of the genomic DNA region transcribed in MCs are summarized in **Fig. 1G**. GATA2 has been identified as a critical transcription factor for the expression of ST2L mRNA, which is driven from the distal promoter in human MCs and basophils^1^. Therefore, we investigated the role of GATA2 in the expression of sST2. As shown in **Fig. 2A**, sST2 mRNA levels decreased in *GATA2* siRNA-transfected LAD2. In contrast, the knockdown (KD) of *GATA1* did not suppress the transcription of sST2 in LAD2. The decrease induced in sST2 mRNA by *GATA2* KD was reflected by a reduction in the release of the sST2 protein from stimulated LAD2 (**Fig. 2B**). The effects of GATA2, but not GATA1, on sST2 mRNA levels of were also observed in the KD experiment using human primary MCs (**Fig. 2C**). Furthermore, sST2 mRNA levels significantly decreased in *Gata2*-KD BMMCs (**Fig. 2D**). These results indicate the important role of GATA2 in sST2 expression in mouse and human MCs.

**Figure 1.**
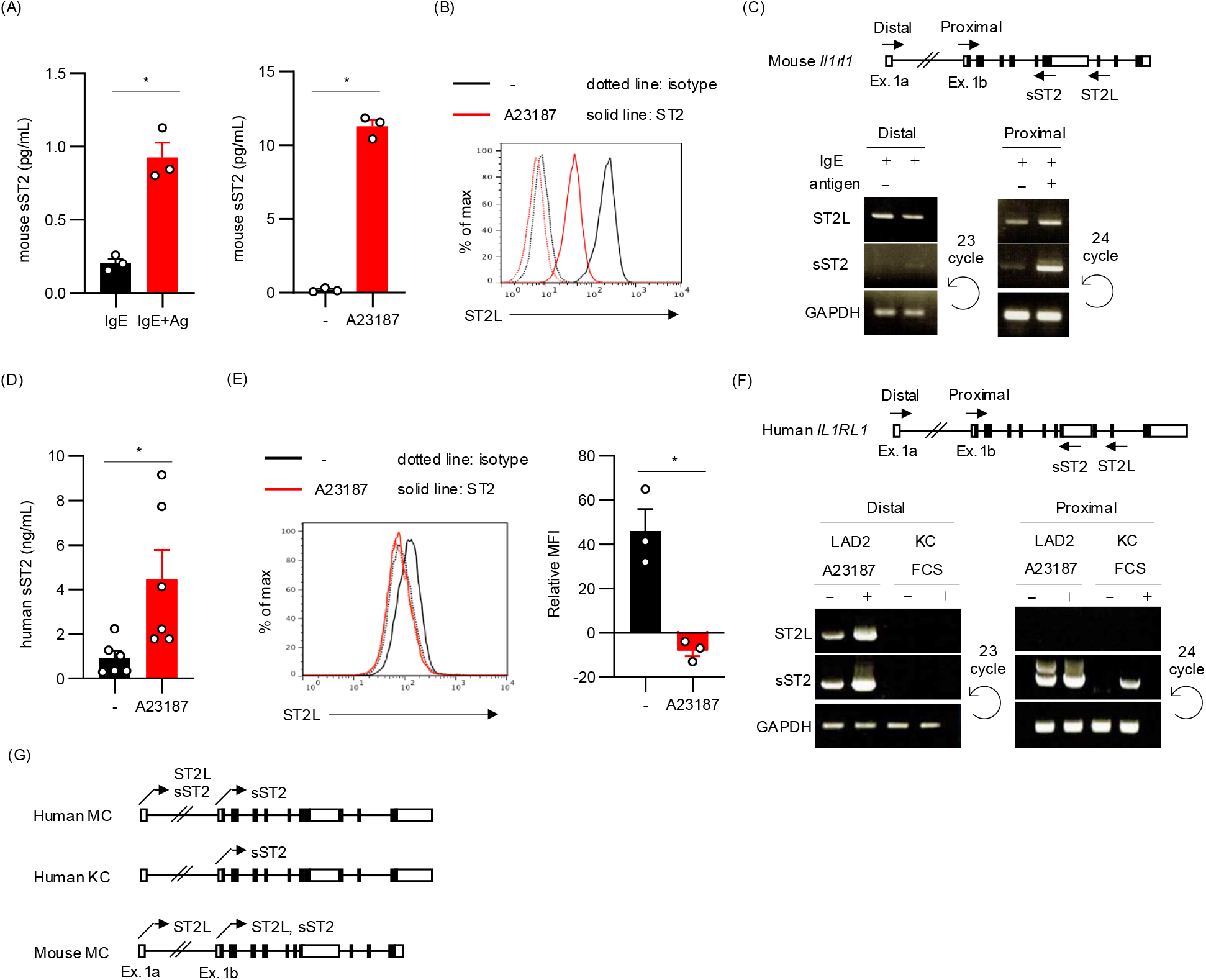
mRNA and protein expression levels of sST2 and ST2L in mouse and human MCs. **(A)** Concentrations of mouse sST2 in culture supernatants of BMMCs. BMMCs (5.0 × 10^5^/μL) were incubated with or without A23187 (left) or IgE plus an antigen (right) for 20 h and 8 h, respectively. **(B)** Cell surface levels of ST2L on BMMCs that were treated with A23187 for 20 h. **(C)** Agarose gel electrophoresis profiles of PCR products (right) amplified using primers designed on the mouse *Il1rl1* gene (left). BMMCs treated with 0.2 μg/mL IgE for 2 h on ice were washed out and then further incubated with (+) or without (-) antigen for 30 min at 37 °C. **(D)** Concentration of human sST2 in culture supernatants of LAD2 (5.0 × 10^5^/μL) incubated with or without A23187. **(E)** Cell surface levels of ST2L on LAD2 treated with A23187 for 16 h. Representative results are shown on the left, and the means ± SEM of three independent experiments are shown on the right. (The relative MFI of ST2L) = (the MFI of ST2L) / (the MFI of isotype control) **(F)** Agarose gel electrophoresis profiles of PCR products (right) amplified using primers designed on the human *IL1RL1* gene (left). LAD2 incubated with (+) or without (-) A23187 and human keratinocyte cells (KC) treated with (+) or without (-) fetal calf serum. A two-tailed Student’s *t*-test was used to compare two samples in Fig. **1A, D**, and **E**. *, *p* < 0.05. **(G)** Genomic DNA regions of the human and mouse *IL1RL1* genes transcribed in MCs.

**Figure 2.**
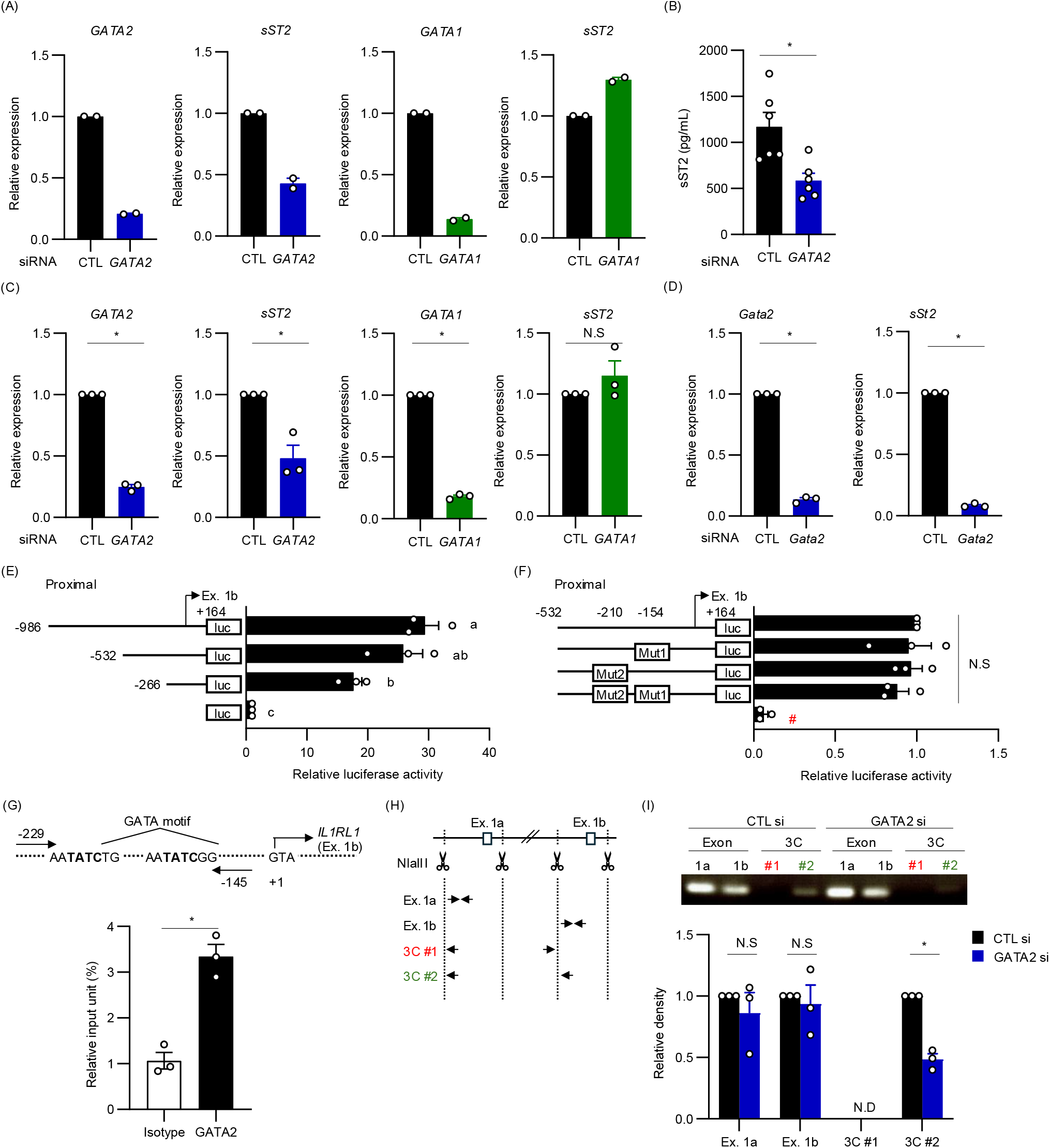
Role of GATA2 in sST2 expression in human MCs. **(A)** mRNA levels of *sST2, GATA2*, and *GATA1* in siRNA-transfected LAD2. **(B)** Concentrations of human sST2 in culture supernatants of LAD2. LAD2 cells (5.0 × 10^5^/μL), which were transfected with control siRNA or *GATA2* siRNA, were incubated for 48 h. **(C)** mRNA levels of *sST2, GATA2*, and *GATA1* in siRNA-transfected primary human MCs generated from peripheral blood CD34^+^ cells in our previous study ^5^. **(D)** mRNA levels of *sST2, ST2L*, and *Gata2* in siRNA-transfected BMMCs. **(E)** A luciferase assay to identify the promoter region of the *IL1RL1* gene. Reporter plasmids carrying various lengths of the 5’-flanking region of the *IL1RL1* gene were transfected into LAD2 cells. Relative luciferase activity was shown as the ratio to that with an empty plasmid. **(F)** Luciferase activities driven by the minimum proximal promoter and its GATA mutants. **(G)** ChIP assay data showing GATA2 binding to the proximal promoter of human *IL1RL1* in LAD2. IgG; isotype control, GATA2; anti-GATA2 Ab. **(H)** Schematic drawing of primers designed for the 3C assay. **(I)** The amounts of PCR products amplified from genomic DNA prepared from *GATA2* KD or its control LAD2 cells. Representative results are shown in the top, and the means ± SEM of relative band intensities in three independent experiments are shown in the bottom. CTL si; Control siRNA-transfected LAD2, GATA2 si; *GATA2* siRNA-transfected LAD2. Data represent the mean ± SEM of two (**A**) or three (**B, C, D, E, F, G, I**) independent experiments. The two-tailed paired *t*-test (**B, C, D, G, I**) and Tukey’s multiple comparison test (**E, F**) were used for statistical analyses. *; *p* < 0.05, N.S.; not significant, N.D.; not detected.

The proximal promoter, which was identified through a reporter assay using deletion constructs (**Fig. 2E**), exhibited similar activity to that of intact constructs even when GATA sequences were mutated (**Fig. 2F**). In contrast, a ChIP assay revealed that GATA2 significantly bound around a minimized proximal promoter region (**Fig. 2G**). Based on these results, we hypothesized that GATA2 transactivates the proximal promoter by binding to the GATA motif(s) in other region(s), including the distal promoter (**Fig. 2H**). This hypothesis was supported by the results of the chromosome conformation capture (3C) assay showing that the amount of genomic DNA amplified by a primer set (a forward primer designed on the proximal promoter and a reverse primer on the proximal promoter) was significantly decreased by *GATA2* KD, while *GATA2* KD had no effect on amplification with the primer set for the distal or proximal promoter (**Fig. 2I**).

Therefore, GATA2 plays a critical role in the expression of sST2 in MCs, which may contribute to the higher levels of sST2 released from MCs than from other cell types. A recent study reported that the production of sST2 from MCs was enhanced by PGE_2_, which ameliorated asthma ^3^. We found that butyrate suppressed IgE-dependent anaphylaxis by accelerating the release of PGE_2_ from MCs ^4^. These findings indicate that the regulation of sST2 production from MCs is a therapeutic target for the prevention of type 2 allergic diseases.

## Authorship Contribution

Contribution: K.N. performed experiments, analyzed data, and prepared figures; K.K. designed the research, performed experiments, and analyzed data; N.I. and K.I. performed experiments and analyzed data; M.H. and K.O. provided experimental tools; C.N. designed the research and wrote the manuscript.

## Acknowledgments

We thank the members of the Laboratory of Molecular Biology and Immunology, Department of Biological Science and Technology, Tokyo University of Science for their constructive discussions and technical support. We greatly appreciate the consideration and support of Dr. Kimihiko Yasuda, Dr. Masako Yasuda, and the late Ms. Yayoi Yasuda.

## Funding Information

This work was supported by Grants-in-Aid for Scientific Research (B) 23K26860 (CN), 23H02167 (CN) and 20H02939 (CN); a Research Fellowship for Young Scientists DC2 and a Grant-in-Aid for JSPS Fellows 21J12113 (KN); a Scholarship for a Doctoral Student in Immunology (from Japanese Society for Immunology to NI); a Tokyo University of Science Grant for President’s Research Promotion (CN); the Tojuro Iijima Foundation for Food Science and Technology (CN); a Research Grant from the Mishima Kaiun Memorial Foundation (CN); and a Research Grant from the Takeda Science Foundation (CN).

## Conflict of Interest Statement

The authors declare no conflict of interest in relation to this work.

## Data Availability Statement

Data that support the present results are available from the corresponding author upon reasonable request.

## Supporting Information

Supplementary Material for this article may be found online.

## Notes

### Competing Interest Statement

The authors have declared no competing interest.

