## Supplemental Information for "The role of GATA2 in the expression of the soluble decoy receptor ST2/IL1RL1 in human and mouse mast cells"

**Supplemental Materials and Methods**

***Cells and Mice***

Bone marrow-derived mast cells (BMMCs) were prepared from C57BL/6 mice (Japan SLC, Hamamatsu, Japan) as previously described ^1^. Animal experiments were performed in accordance with the approved guidelines of the Institutional Review Board of Tokyo University of Science, and the Animal Care and Use Committees of Tokyo University of Science approved this study. The present study was approved by the Animal Care and Use Committees of Tokyo University of Science: K22005, K21004, K20005, K19006, K18006, K17009, and K17012. The LAD2 (human mast cell leukemia) was kindly provided by Dr. Arnold Kirshenbaum and maintained in the human SCF-supplemented medium ^2^. Human primary mast cell data was obtained by re-analyzing the samples obtained in our previous study ^3^. The stimulation of BMMCs with IgE (anti-TNP mouse IgE clone IgE-3, BD Bioscience, San Jose, CA) and antigen (TNP-BSA, LSL, Tokyo, Japan) was performed as previously described ^4^. Calcium ionophore A23187 (CAS RN: 52665-69-7), and recombinant mouse IL-33 (#210-33, Peprotech) were also used to stimulate MCs.

***ELISA***

Concentration of mouse sST2 and human sST2 was determined by Quantikine ELISA kit (R&D Systems, #MST200 and #DST200, respectively).

***Flow cytometry***

Cell surface expression levels of mouse and human ST2L were determined by a MACS Quant Analyzer (Miltenyi Biotech) using FITC-labeled anti-mouse ST2L Ab (mdbioproducts, clone: DJ8) and PE-labeled anti-human ST2L Ab (MBL life science, clone: 2A5), respectively.

***Semi-quantitative PCR of mRNA***

A ReliaPrep RNA cell miniprep system (Promega) and ReverTra Ace qPCR RT Master Mix (TOYOBO) were used to purify total RNA and to synthesized cDNA, respectively. We used following primers for semi-quantitative PCRs.

For the mouse *Il1rl1* gene: distal; 5’- GATGGCTAGGACCTCTGGC-3’, proximal; 5’- GTAGCCTCACGGCTCTGAGCT-3’, ST2L; 5’- AGCAACCTCAATCCAGAACACT-3’, sST2; 5’-TGGAAGACAGAAACATTCTGGA-3’.

For the human *IL1RL1* gene: distal; 5’-GTTGTGAAACTGTGGGCAGA-3’, proximal; 5’- CAAATTTCAGGATGGGAGGA-3’, ST2L; 5’-ATGTCTCTCCAGAGCAGAGTGG-3’, sST2; 5’- GCCATTCCCATTGTCTTGAT-3’.

A common primer set for mouse and human *Gapdh*/*GAPDH*: forward; 5’-TGAACGGGAAGCTCACTGG-3’, reverse; 5’-TCCACCACCCTGTTGC TGTA-3’.

***Quantification of mRNA***

Quantification of mRNAs was performed as previously described ^4^. TaqMan probes and primers are as listed follows.

Human *GATA2*; Hs00231119_m1, human sST2 (Ex 1b); Hs01073297_m1; human *GATA1*; Hs01085823_m1, human *GAPDH*; 4326317E, mouse *Gata2*; Mm00492300_m1, mouse *Gata1*; Mm01352636_m1, mouse *Gapdh*; 4352339E.

A primer set for mouse sST2: forward; CGTCTGCAGAAATGAG, reverse; TATGGCAATGGCACAG.

***Knockdown using small interfering RNA (siRNA)***

Knockdowns of human *Gata2* mRNA in LAD2 cells and mouse *Gata2* mRNA in BMMCs by using siRNAs were performed as previously described ^1, 3, 5^.

***Reporter assay***

A reporter plasmid carrying distal promoter region of the human *IL1RL1* gene has been generated in our previous study ^5^, and proximal promoter region (-532/+164) of the human *IL1RL1* gene amplified by PCR were introduced into pGL-4.10[luc2] (Promega, #E6651) to obtain a luciferase reporter plasmid of the proximal promoter. Transfection of reporter plasmids into LAD2 cells and determination of luciferase activity were performed as previously described ^5^.

***Chromatin immunoprecipitation (ChIP) assay***

ChIP assay was performed as previously described ^1, 5^.

***Chromosome conformation capture (3C) assay***

We performed 3C assay based on the method originally established in a previous study^6^ using following primers.

Ex1a forward; 5’- TTCAAATAGGGAGAATGTGGTGAA-3’, Ex1a reverse; 5’-CTGAACCCGATGATCACACTGT-3’, Ex1b forward; 5’-AAACTCCTGGCACCAATGCT-3’, Ex1b reverse; 5’-TGGCAGACACCACTCAGGAA-3’, 3C primer 1; 5’-TGTCTTAACTATATTTTGCCTTCTTCATG-3’, 3C primer 2; 5’-GTTGATTCTAAAATAGGAGGAAATGATTC-3’, 3C primer 3; 5’-CCTCCCATCCTGAAATTTGATTT-3’.

***Statistical analysis***

A two-tailed Student’s *t*-test was used to compare two samples, and a one-way ANOVA followed by Tukey’s multiple comparison test and Dunnett’s multiple comparison test were performed to compare more than three samples. *P* values < 0.05 were considered to be significant.
